# Variation in gene expression patterns across a conifer hybrid zone highlights the architecture of adaptive evolution under novel selective pressures

**DOI:** 10.1101/2021.11.24.469945

**Authors:** Mitra Menon, Jared Swenson, Ehren Moler, Amy V. Whipple, Kristen M. Waring, Andrew J. Eckert

**Affiliations:** Department of Evolution & Ecology and Center for Population Biology, University of California, Davis, CA 95616, USA; Department of Biology, Virginia Commonwealth University, Richmond, VA 23284, USA; Department of Biological Sciences and Center for Adaptive Western Landscapes, Northern Arizona University, Flagstaff, AZ 86011, USA; School of Forestry, Northern Arizona University, Flagstaff, AZ 86011, USA

## Abstract

- Variation in gene expression among natural populations are key contributors to adaptive evolution. Understanding the architecture underlying adaptive trait evolution provides insights into the adaptive potential of populations exposed to novel selective pressures.
- We investigated patterns and processes driving trait differentiation under novel climatic conditions by combining common garden experiments with transcriptome-wide datasets obtained from *Pinus strobiformis - Pinus flexilis* hybrid zone populations.
- We found strong signals of genotype-environment interactions at the individual transcript and the co-expression module level, a marked influence of drought related variables on adaptive evolution and an environment dependent influence of *P. flexilis* ancestry on survival. Using co-expression network connectivity as a proxy for pleiotropy we highlight that adaptive transcripts were pleiotropic across both gardens and modules with strong population differentiation exhibited lower preservation across gardens.
- Our work highlights the utility of integrating transcriptomics with space-for-time substitution studies to evaluate the adaptive potential of long-lived species. We conclude by suggesting that the combination of pleiotropic trait architectures and substantial genetic variation may enable long-lived forest tree species to respond to rapid shift in climatic conditions.

## Introduction

Spatial variation in selection pressures can lead to ecological specialisation and genetic differentiation among populations. Such ecological specialisation is often referred to as local adaptation and is a key contributor towards biodiversity. Common garden experiments and provenance trials of numerous species have demonstrated local adaptation via a fitness reduction when populations are translocated to abiotic and biotic conditions that diverge from those found in their home environment (Clausen et al. 1948; Savolainen et al. 2013; Pickles et al. 2015). Common garden studies have also revealed among-population variation in phenotypic plasticity (Dayan et al. 2015), defined as the environment-dependent expression of a trait value. Local adaptation and phenotypic plasticity are commonly represented by genotype-by-environment interactions (GEI) and hypothesised as major mechanisms facilitating survival under changing climatic conditions (Bradshaw, 2006; Nicotra et al. 2015). Despite the abundance of studies investigating local adaptation, very few have evaluated GEI and the architecture underlying it under novel selection pressures. Such novel selection pressures can be imposed through space-for-time substitution designs (*sensu* Pickett, 1989) conducted by using common gardens located beyond or at the climate margin of a species range (Geber and Eckhart, 2005). Most empirical studies of adaptive evolution have been in line with Fisher’s theoretical prediction of variation in quantitative traits being driven by many small effect loci (Fisher, 193). But contrary to the relatively higher contribution of small effect loci under migration-selection balance, we might expect a disproportionate effect of large effect loci during initial stages of adaptation to novel selective pressures (Orr, 1998; Hamala et al. 2021).

Studies of forest trees often reveal strong patterns of local adaptation at broad and fine spatial scales (Csilléry et al. 2014; reviewed in Lind et al. 2018). Across most forest trees and in general for species exhibiting limited isolating barriers, both intra- and inter-specific gene flow are key sources of adaptive evolution, specifically when faced with novel selective pressures (Aitken et al. 2008; Taylor & Larson, 2019; Bolte & Eckert, 2020). This is likely because inter-specific gene flow via hybridization increases additive genetic variation, which influences trait responses to selection (Falconer & McKay, 1996). Studies in spruce, pine and poplar that utilize fitness-associated quantitative traits such as height, volume, bud set and disease resistance have demonstrated higher heritability and phenotypic variance in hybrid populations compared to non-hybridized populations, and better hybrid performance in novel environmental conditions (reviewed in Dungey, 2001; Suarez-Gonzalez et al. 2016, 2018; De La Torre et al. 2014). However, investigations of the architecture underlying GEI under novel environments and the contribution of hybrid ancestry towards GEI have lagged behind traditional investigations of adaptive evolution in forest trees.

Combining global gene expression datasets with survival – a key fitness component-can help overcome the challenge posed by longevity of trees. Gene expression variation in natural populations is often highly heritable, is influenced by genetic and environmental variation, and thus reflects signatures of various selective pressures (Whitehead & Crawford, 2006). By capitalizing on the modular nature of gene expression patterns (Hartwell et al. 1999) and treating gene expression as a quantitative trait (*see* Roberge et al. 2007), we can more thoroughly evaluate the architecture that may be favoured under novel selective pressures. Data obtained from transcriptomic profiling can be organized into co-expression modules. This approach relies on connectivity patterns among genes and assumes that strongly connected genes are reflective of similar functional categories and exhibit coordinated response to selective pressures. Connectivity patterns within a co-expression module can be shaped by selection, such that highly connected “core” genes are often pleiotropic and experience stronger selective constraints, while “peripheral” genes with lower connectivity are often involved in GEI (Cork & Purugganan, 2004; Josephs et al. 2017).

Our study utilizes the natural hybrid zone formed between two ecologically divergent species of pine, *Pinus strobiformis* Engelm. and *P. flexilis* E. James, to evaluate signatures of GEI. Both species have broad geographic distributions across western North America, with hybrid populations inhabiting fragmented sky-island ecosystems of New Mexico, Arizona, Texas and southern Colorado (Bisbee, 2014; Menon et al. 2018) where they experience ongoing gene flow from *P. flexilis* (Menon et al. 2018; Critchfield,1975). These hybrid populations experience severe drought, aseasonal frost and diurnal fluctuations in temperature (Adams & Kolb, 2004; Williams et al. 2010). In agreement with theoretical models (Fisher, 1930), previous genotype-environment association studies in this system have demonstrated a dominance of many small effect loci driving adaptive evolution (Menon et al. 2021). However, an understanding of how adaptive evolution proceeds under novel selective pressures is only beginning to be gathered in trees and across other systems.

The overarching goal of this work was to evaluate the interaction between interspecific gene flow and novel selective pressures on the genomic architecture of GEI. To simulate novel selective pressures, we utilized a space-for-time substitution design wherein we planted the hybrid seedlings across two common gardens that represent the extremes of their suitable habitat. One garden represented a warmer, more arid climate while the second garden represented a combination of higher aridity and increased freeze events (Fig. S1). Together our design is representative of the global increase in aridity and warming along with shifts in multivariate environmental axes as is projected to occur in the western US under all plausible climate change scenarios (Cook et al. 2015). By coupling our hybrid zone populations with a space-for-time substitution design assaying gene expression patterns we test the following hypotheses. **(H1)** Our assayed population will demonstrate strong signals of adaptive trait differentiation. This will reflect the heterogeneity in source population environmental conditions as well as the novel selective pressures that our seedlings were exposed to. **(H2)** Differences in the selective environment between the two gardens will result in garden specific patterns of trait differentiation. However, given the potential physiological and cellular cross-talk between various stress tolerance strategies (Wong et al. 2005), we expect to note some degree of shared patterns of differentiation. As a result, while overall co-expression network preservation may be strong, modules exhibiting BS specific population differentiation may have generally low preservation in WP and vice-versa. **(H3)** Hybrid genomic ancestry will impact GEI. Drawing from our previous work demonstrating adaptive introgression of *P. flexilis-like* loci only along freeze-related environmental gradients, we expect higher contribution of ancestry towards adaptive evolution and towards increased survival in the colder garden. **(H4)** The architecture of adaptive evolution will be dominated by weakly connected “peripheral” genes in the co-expression modules given that these are less likely to have negative pleiotropic consequences and hence amenable to be fine-tuned in response to changing selective pressures.

## Methods & Materials

### Common garden design

This study makes use of seedlings obtained from 30 maternal trees that were replicated across two common gardens established in the Kaibab National Forest, AZ (Fig. 1a & b) as part of the Southwest Experimental Garden Array (SEGA). Seedlings within each garden were planted in a randomized block design with siblings representing members of each of the 30 maternal families (Squillace, 1974). In August 2019, we sampled 90 seedlings per garden (3 siblings/maternal family) representing 30 maternal families originating from 10 populations (3 maternal trees/population) (Fig. 1b) sampled across the *Pinus strobiformis-P. flexilis* hybrid zone (Menon et al. 2018). Our sampling year and design provided the opportunity to assess how trees will respond to novel selective pressures. During 2019, the high elevation garden, Bear Springs (BS), experienced drier and colder conditions relative to the values noted across a 60-year period in the native range of the sampled hybrid populations. The low elevation garden, White Pockets (WP), experienced warmer and drier conditions relative to the sampled populations and to BS as well (Fig. S1). The seedlings were further exposed to a 50% reduction in ambient precipitation for 3 growing seasons prior to our sampling in 2019.

**Figure 1:**
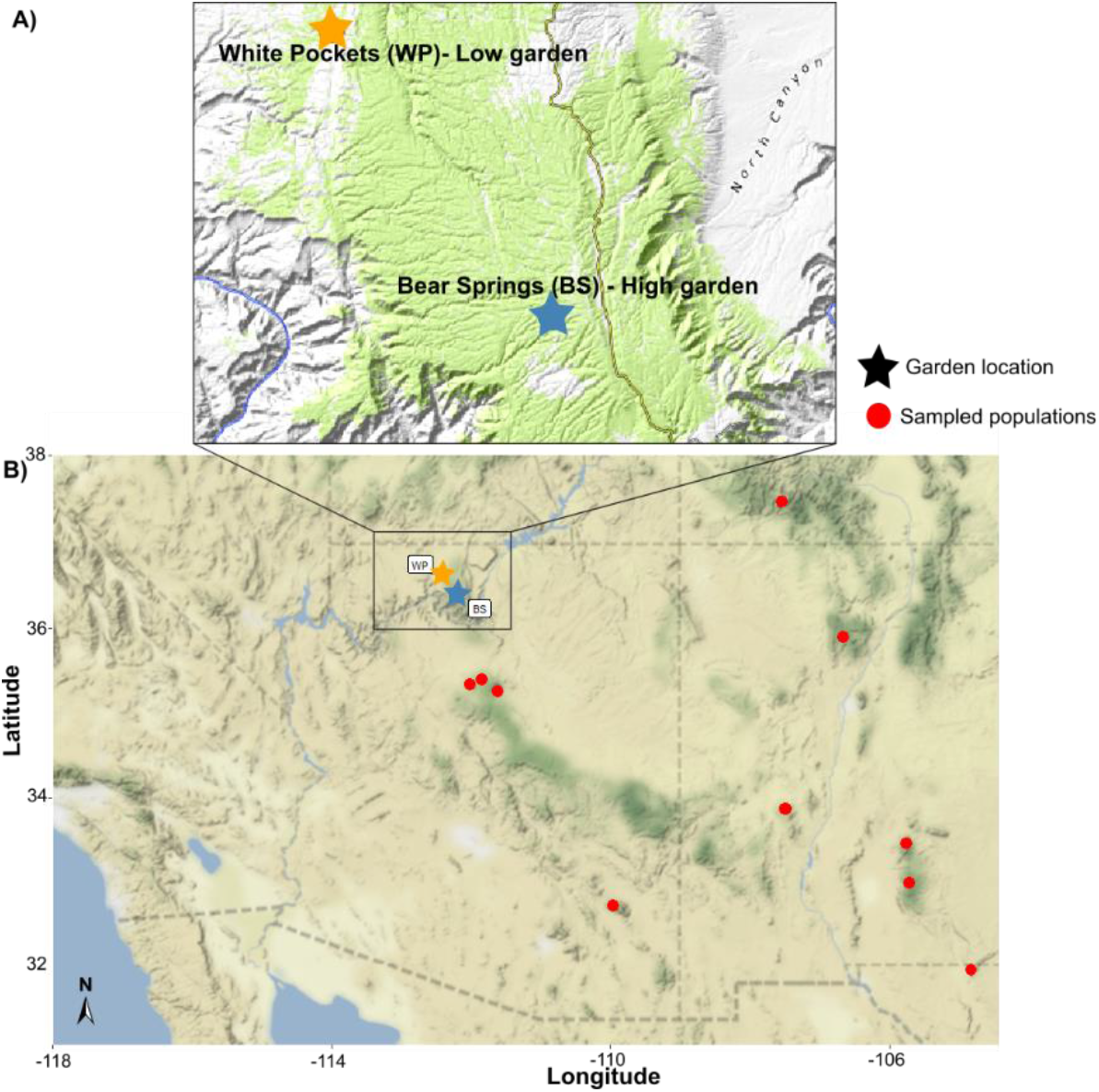
a) Location of the two common gardens on the Kaibab National Forest and b) Location of the sampled populations in relation to the common garden sites.

### Transcriptomic dataset

For each seedling, a maximum of three fascicles from the current year’s growth were sampled and flash frozen in liquid nitrogen for RNA extraction. Sampling was done across two days with consistent weather conditions between the hours of 12:00 and 17:00. A maximum of 100 mg of needle tissue per seedling was ground into a fine powder using liquid nitrogen with a mortar and pestle by adding in 40 mg of Polyvinylpyrrolidone (PVP-40). Total RNA was extracted using Spectrum Plant total RNA kit (Sigma-Aldrich) with slight modifications to the manufacturer’s protocol. Following polyA tail selection to enrich for mRNA, paired-end (2 × 150bp) RNA libraries were prepared following standard protocols for NEBNext Ultra II RNA Library Prep (Illumina) and sequenced on the Novaseq6000 platform at Novogene Co. (Sacramento, CA).

Raw fastq files were assessed for quality using fastqc and trimmed to remove adapters and low quality reads using TrimGalore v.0.6.4 (Krueger, 2015). Due to lack of a well annotated genome from a closely related species, we built a *de novo* transcriptome assembly using Trinity v.2.8.5 (Grabherr et al. 2011; Haas et al. 2013) with a *k*-mer length of 24 and a minimum contig size of 300bp. Further details about the assembly pipeline, filtering steps, quality assessment and annotations using EnTAP (Hart et al. 2020) are detailed in the Supporting Information (Methods S1).

### Genomic dataset & estimation of hybrid ancestry

To evaluate genome-wide divergence among the populations used in our transcriptome sampling and to estimate the influence of hybrid ancestry on gene expression patterns we obtained genomic DNA from 270 trees using DNeasy 96 Plant Kit (Qiagen). These 270 trees included the 30 maternal trees represented in our mixed-sib family design along with previously identified 154 pure *P. strobiformis* and 86 pure *P. flexilis* individuals (Menon et al. 2018; Fig. S2a). Following the ddRADseq library preparation detailed in Parchman et al. (2013), we size selected 300-400 bp regions and performed single-end sequencing (1 × 150 bp) on Illumina HiSeq 4000 (Novogene Corporation). The resulting fastq files were processed using dDocent v.1.0 (Puritz et al. 2014) to obtain genotype calls, as in Menon et al. (2018, 2021). Following post-filtering steps implemented in VCFTOOLS v.0.1.153 (Danecek et al. 2011) and custom functions implemented through python v.3 (Methods S2), we obtained 23,261 SNPs. We used this full dataset of 270 individuals and 23,261 SNPs to estimate hybrid ancestry through the genotype likelihood-based clustering algorithm implemented in NGSAdmix (Skotte et al. 2013) with *K*=2 representing the two hybridizing species.

### Estimation of maternal expression trait values

Prior to conducting further analyses using the 47,651 transcripts obtained after a series of filtering steps (Methods S1), we normalised the read count data for varying library sizes using the Trimmed mean of M values (TMM) approach implemented within the calcNormFactors function from edgeR v.3.14.0 (Robinson et al. 2010) in R v.4.0.2 (R Core Team, 2020). We then filtered out transcripts that were not expressed across all three sibs nested within the same maternal family in either garden. To evaluate signatures of adaptive trait differentiation we implemented a mixed effects model (eq. 1) that utilized the sib design and determined each maternal family’s normalised expression value for the 27,441 transcripts retained from our filtering steps:

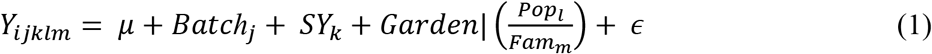

Here, *Y* represents the normalised expression of the transcript for seedling *i*, *μ* represents the global mean for the transcript. *Batch* and *SY* are fixed effects and represent the date of sequencing and year of planting, respectively. *Fam* represents the maternal family which is nested within *Pop* (representing the source population of the maternal family) and treated as the random effect which is allowed to vary by *Garden*. This linear mixed effect model was fitted using the *fitVarPartModel()* function in the variancePartition v.1.16.1 package (Hoffman & Schadt, 2016) in R, which uses log-transformed normalised counts and precision weights obtained from *voomWithDreamWeights*. Precision weights are ideal to model mean-variance trends typical of RNAseq datasets (Law et al. 2014). Using the fitted model in (1), we then obtained garden-specific maternal values and garden-specific population values for each transcript.

### Assessing per-transcript level signatures of local adaptation

We treated expression as a quantitative trait (see Lasky et al. 2014; Huang et al. 2020) and evaluated the among population component of heritable variance (*Q_ST_*) (Spitze, 1993). Under neutrality, we expect *Q_ST_* to be similar to its genome wide equivalent (*F_ST_*) (Leinonen et al. 2008), but under spatially divergent selection pressures driving local adaptation *Q*_ST_ should exceed *F*_ST_. To generate our expectation of differentiation under neutrality, we retained 11,432 biallelic SNPs identified across only the 30 maternal families for which gene expression data were available. For these SNPs we estimated *F_ST_* using the method of Weir and Cockerham (1984) as implemented in the HIERFSTAT package v.0.04-22 in R (Goudet, 2005). Using the per transcript variance components obtained from equation (1) we estimated *Q_ST_* as:

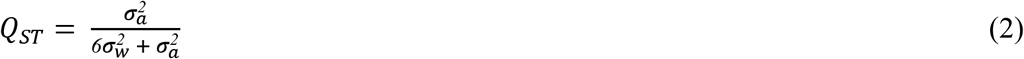

where 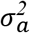 represents the among population variance component and 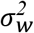 represents the within population (among family) variance components. The constant 6 represents the mixed-sib design assuming a 50:50 mixture of half-sibs and full-sibs. Confidence intervals (CI, 95%) per transcript per garden were obtained using 1000 parametric bootstrap estimates generated using the *bootMer*() function implemented in lme4 (Bates et al. 2015). To address hypothesis H1 at the per trait level we compared the 0.025 quantile of *Q_ST_* for each transcript with the 0.95 quantile of *F_ST_* and identified transcripts exhibiting signatures of adaptive expression differentiation across gardens and unique to each garden. Since we are primarily interested in the architecture underlying GEI, subsequent analyses and our main results focus only on three *Q_ST_* categories: (a) conditionally adaptive in BS (BS-condA), transcripts with *Q_ST_* > *F_ST_* only in high elevation garden; (b) conditionally adaptive in WP (WP-condA), transcripts with *Q_ST_* > *F_ST_* only in low elevation garden; and (c) adaptive plasticity (Ad-Pl), transcripts with *Q_ST_* > *F_ST_* across both gardens and a significant *Q_ST_* reaction norm (*p* < 0.05) assessed through a t-test (Fig. 2). For each of these *Q_ST_* categories we performed gene ontology (GO) enrichment analyses using a hypergeometric test with 1000 permutations to compute the family-wise error rate as implemented in the GOfuncR v1.12 package (Grote, 2020) in R. The background for GO enrichment analyses was the full set of GO terms across all annotated and non-contaminant transcripts (Table S1).

**Figure 2:**
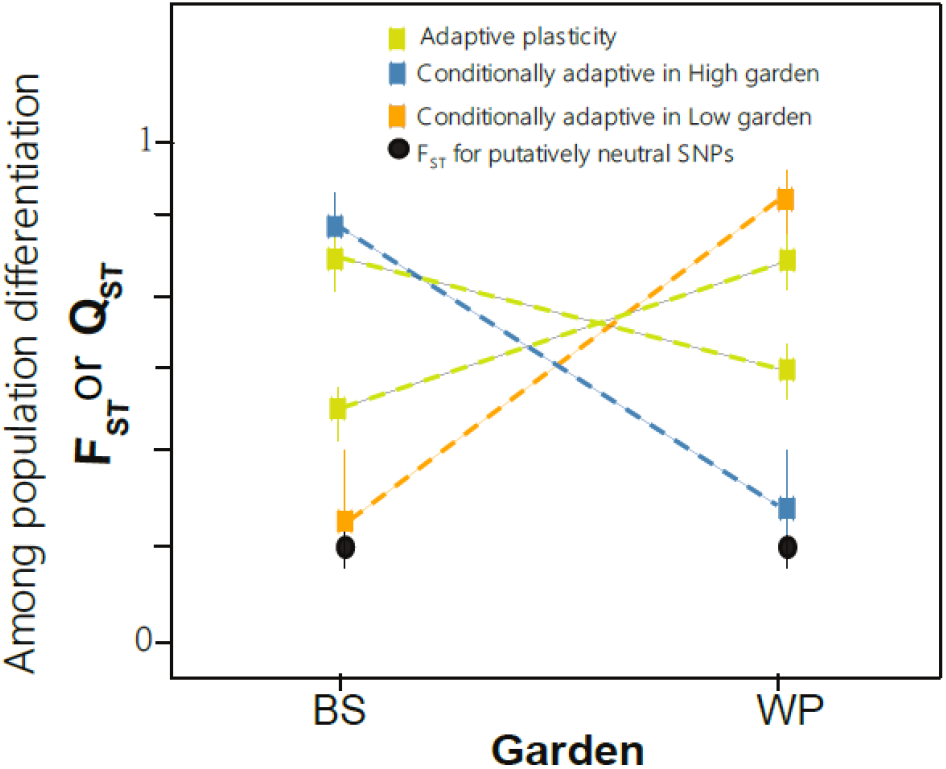
Schematic representation of *Q_ST_* categories used in this study. Numbers next to each category indicate the number of transcripts classified under them using the *Q_ST_* - *F_ST_* comparison.

### Assessing co-expression module level signatures of local adaptation

Local adaptation is often influenced by trait covariation (O’Brien et al. 2019), thus modules formed by sets of strongly correlated transcripts can be effectively used to capture signatures of local adaptation. We made use of modules identified from gene co-expression networks constructed per garden using the WGCNA v1.70 package (Langdelfer & Horvath, 2008) in R. Since WGCNA does not implicitly account for the study design, we used the family level transcript abundance estimated from eq. 1 as input. Thus, in our case the co-expression network represents sets of transcripts with correlated expression levels across maternal families in a given garden. Modules formed within each co-expression network were visualised using the prefuse force directed edge weighted layout in Cytoscape v.3.8.2 (Shannon et al. 2003). To highlight transcripts of interest and reduce complexity we filtered out weakly connected transcripts and edges with weights below the 75th quantile for the module (Fig. S3). Transcripts falling within each module were used to perform GO enrichment analyses using the same approach as described above for the three *Q*_ST_ categories of interest. To address hypothesis H1 at the functional module level we evaluated among population differences in the co-expression module using a linear model with the first eigengene of each module as the response variable and population of origin as the predictor variable. We also assessed the relative representation of individual adaptive transcripts (identified using *Q*_ST_- *F*_ST_ approach above, Fig 2) in each module (eq. 3). For modules detected at BS, we used the BS-condA as the *Q*_ST_ category, while for modules in detected in WP we used the WP-condA category. In all cases, relative representation was estimated as:

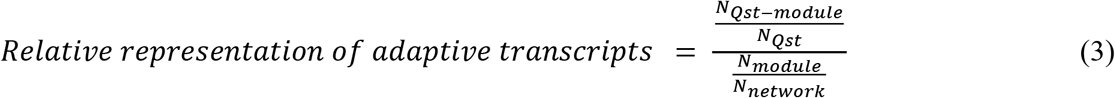

where N_*Q*ST -module_ is the number of transcripts in a module that were also classified under the respective *Q*_ST_ category, N_*Q*ST_ is the number of transcripts identified under the *Q*_ST_ category (Fig. 2), N_module_ is the total number of transcripts in a module, and N_network_ is the total number of transcripts used in WGCNA. Statistically significant over- or under-represented modules (two tailed p < 0.05) were identified by permuting the transcript labels and calculating 10,000 null relative representation estimates for each module.

### Estimating the effect of environment on among population differentiation

To further address hypothesis H1 and H2 at the per-transcript level we identified climatic drivers of adaptive differentiation for our three *Q*_ST_ categories (Fig. 2) using redundancy analysis (RDA) in R (vegan package v.2.5.6: Oksanen et al. 2013). Annual and seasonal climate variables at 1 km resolution were obtained from ClimateWNA v6.1 (Wang et al. 2016) for the period 1981-2010 for each of the 30 maternal trees. Population-level estimates of climate variables were obtained by averaging values across the three maternal trees selected per population. Following Menon et al. (2021), we classified the environmental variables into drought- and freeze-associated variables, as these two axes are likely the primary selective agents across the hybrid zone populations. To reduce the dimensionality of the environmental dataset and avoid multicollinearity, we performed principal component analysis (PCA) separately for the drought and freeze associated variables and retained the PC axes that explained at least 90% of the variance as predictors in RDA. We used the population-level estimates of gene expression from eq. (1) as the response matrix with the drought-associated PCs, freeze-associated PCs, and geography as the predictor matrices. Geography was represented by scaled and centered estimates of latitude and longitude. For each *Q*_ST_ category, we evaluated whether among population gene expression differences were primarily impacted by a) all three predictor matrices jointly, b) by the drought-associated matrix, or c) by freeze-associated matrix.

### Influence of P. flexilis ancestry on expression patterns and survival

Survival for all mixed-sibs in the gardens was recorded in August 2019 as a binary trait. To obtain family and population level estimates of survival we fitted a similar mixed effect model as used for expression traits but with binomial error distribution (Swenson et al. *in prep*) implemented with *glmmTMB* (Brooks et al. 2017) in R.

To assess hypothesis H3 at the per transcript level, we correlated mean population-level estimates of *P. flexilis* ancestry obtained from NGSAdmix (Fig. S2b) with population-level expression values for each *Q*_ST_ category. For BS-condA and WP-condA, we used expression values estimated at BS and WP, respectively. For Ad-Pl we used the absolute difference in expression values across gardens. For each *Q*_ST_ category the observed distribution of correlation coefficients was compared against an empirical background distribution that was matched on expression bins to the category of interest. Significant deviation in the observed dataset was evaluated using the Kolmogorov-Smirnov test implemented in R at a threshold of 0.05. Since our results were not sensitive to the binning, and expression levels did not vary much by category (results not shown), we report results only from the simple permutation test. Using a similar approach, we evaluated whether the three *Q*_ST_ categories deviated significantly from the background set of transcripts in their association with the estimated population-level survival measured at the respective gardens (Swenson et al. in prep). For the Ad-Pl category we used the mean of the estimated survival values across gardens. Similarly, we tested H3 at the co-expression module level by evaluating whether the eigengene expression for a module was associated with *P. flexilis* ancestry and with survival using Pearson’s correlation coefficient with multiple testing correction (q < 0.05).

### Connectivity patterns of Q_ST_ transcripts and preservation of modules across gardens

Network topological measures such as gene-gene connectivity and centrality of a gene are often reflective of types and strength of selection acting on the gene (Jordan et al. 2004; Josephs et al. 2017; Des Marais et al. 2017). We obtained three measures of connectivity for each transcript. These reflected intramodular connectivity (kWithin), connectivity to all transcripts disregarding module membership in the network (kTotal) and the difference between inter- and intra-modular connectivity (kDiff).

To evaluate hypothesis H2 from a co-expression network perspective we performed comparative network analyses using 10,000 permutations with the WGCNA package (Langfelder et al. 2011) in R. We utilised two aggregate summary statistics implemented within WGCNA to declare a module as being preserved or not across gardens. *Z_summary_* represents a normalised value of various connectivity and density-based measures following a permutation test procedure, while *medianRank* simply scores each module based on the observed preservation statistic. Following Langfelder et al. (2011), we used both *Z_summary_* and *medianRank* for inference, because while the former is strongly influenced by module size the latter is not. We declared a module as preserved across gardens if its *Z_summary_* score was higher than 10 and *medianRank* below 5. Modules with *Z_summary_* score below 10 and a *medianRank* above 5 had weaker evidence for preservation (Langfelder et al. 2011).

To assess hypothesis H4, we implemented the network characterization approach developed by Mahler et al. (2017) to define the core of a module, while the core of a network was defined as the top 10% of transcripts with highest kTotal. We determined if the observed relationship between *Q_ST_* categories and connectivity was significantly different than expected by chance by performing 10,000 null permutations. Since expression levels are often associated with connectivity in a co-expression network (Fig. S4), our permuted sets were matched on bins of gene expression to be representative of the core transcripts.

## RESULTS

### Transcript and module level signatures of adaptive evolution (H1)

At the per transcript level estimates of *Q_ST_* across both gardens ranged from 0 to 1 (Fig. 2) with a mean of 0.31 ± 0.37 (sd) for BS and 0.37 ± 0.38 for WP. Approximately 50% of the transcripts exhibited differentiation below 0.1 in BS and 40% were below 0.1 in the WP. Only 20% of the transcripts had strong differentiation (above 0.9) in either garden. The distribution of *F_ST_* using genome-wide ddRADseq markers was centered on zero with a multilocus *F_ST_* of 0.015 (Fig. S2.c). Using *Q_ST_-F_ST_* comparisons, we identified 334 transcripts as adaptive in BS, 469 as adaptive in WP, 234 as BS-condA, 344 as WP-condA and 26 transcripts under the category Ad-Pl (Fig. 2).

At BS, our co-expression network consisted of 22,757 transcripts, which were grouped into 31 modules (Methods S3, Fig S5) with the number of transcripts per module ranging from 75 to 5668. At WP, our co-expression network consisted of 23,468 transcripts, which were grouped into 26 modules with the number of transcripts per module ranging from 96 to 9410 (Methods S3, Fig S5). Using the eigengene values of each module we identified 20 modules in WP and 21 in BS exhibiting strong signals of among population differentiation (Table S3). Of these 20 modules identified in WP, five were also significantly enriched for transcripts categorised under WP-condA (p < 0.05). The five modules exhibiting significant depletion for WP-condA transcripts also had weak eigengene level population differentiation (Fig. 4a,b, Table S3). Similarly in BS of the 21 modules, four were also significantly enriched (p < 0.05) for transcripts categorised under BS-condA and three modules that were significantly depleted (p < 0.05) did not exhibit strong module eigengene level population differentiation (Fig. 4c,d, Table S3). In WP, the number of transcripts in enriched modules ranged from 125 to 3439 while in BS they ranged from 139 to 5668. Overall, using a Kruskal-Wallis test, *Q*_ST_ enrichment or depletion was not associated with module size across either garden (BSdf=2 = 4.78, p = 0.09, WPdf=2 = 0.68, p = 0.71).

**Figure 3:**
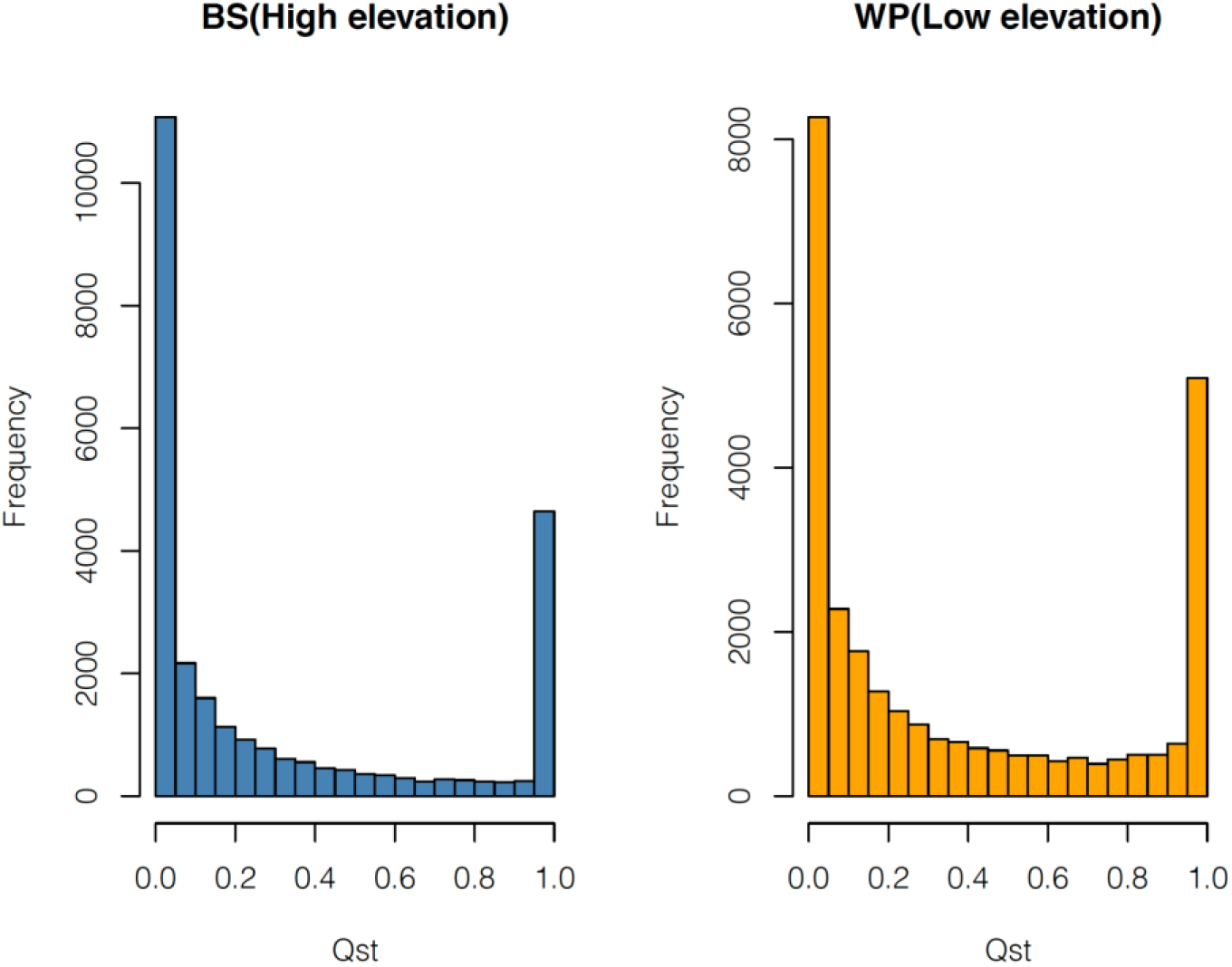
Distribution of transcript expression *Q_ST_* values across gardens.

**Figure 4:**
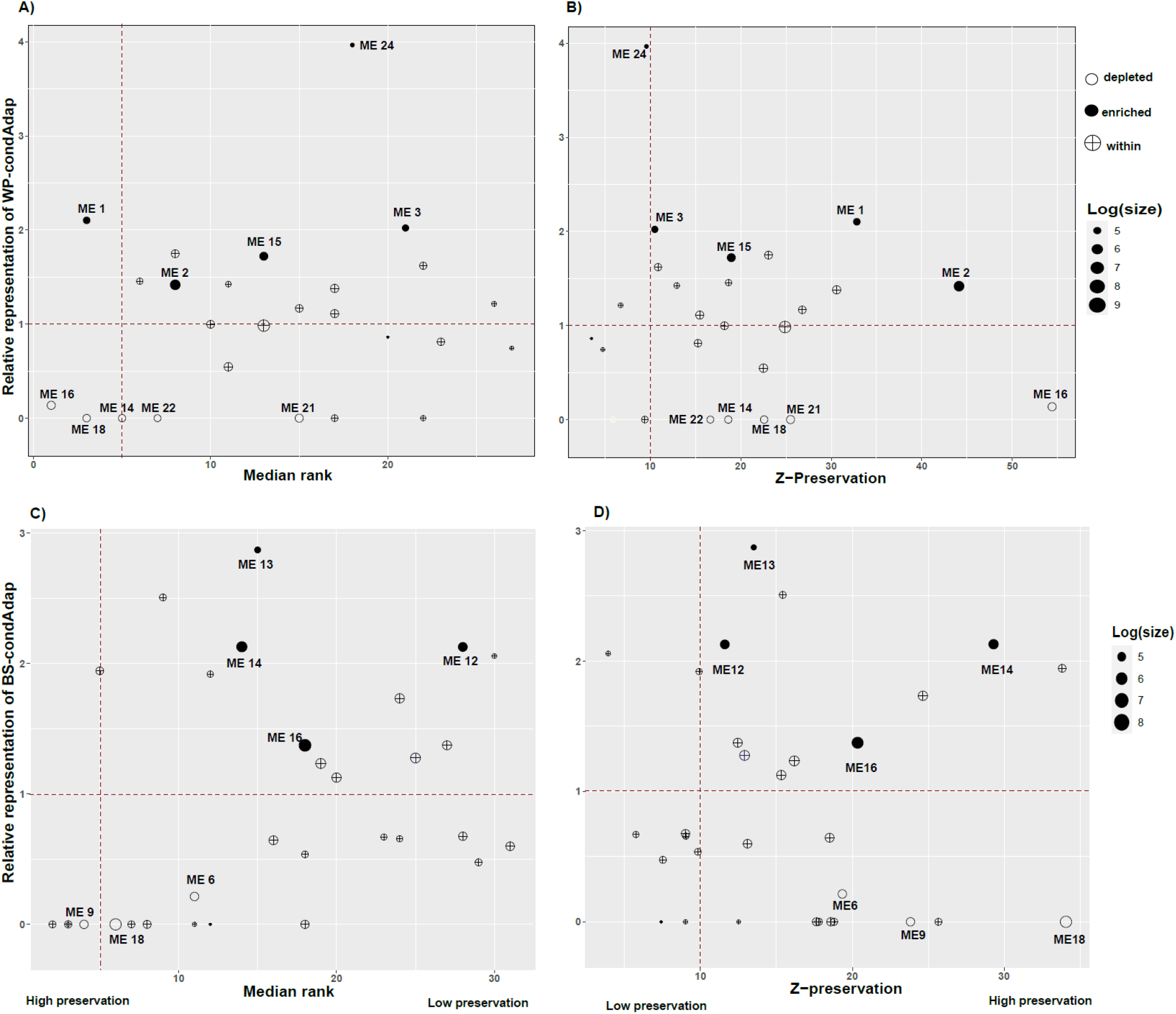
Relationship between *Q*_ST_ transcript enrichment, module size and module preservation measures across gardens for modules identified in **a & b)** WP: Low elevation garden and in **c & d)** BS: High elevation garden. Vertical dotted line indicates the cutoff for declaring a module as preserved using median rank and Z-preservation score. Note that a value below 5 for median rank indicates strongly preserved modules, while modules with Z-preservation score above 10 are preserved. Only modules significantly over-enriched or under-enriched for *Q*_ST_ outlier transcripts have been labelled.

### Drivers of adaptive evolution at the per-transcript level (H2 & H3)

For our three *Q*_ST_ categories of interest (WP-condA, BS-condA and Ad-Pl, see Fig. 2), we evaluated the impact of environmental factors on population level expression differences using RDA. To determine if these transcripts were related to garden specific survival and whether they were influenced by genomic ancestry, we utilised a permutation-based correlation approach.

The full RDA model (model 1) was significant for all three *Q*_ST_ categories and explained 50% of the expression variance for BS-condA category, 17% for the WP-condA category and 19% for Ad-Pl (Table 1). Similarly, the RDA model used to assess the conditional impact of drought (model 2) was significant for all categories and explained 48% of the variance for the BS-condA category, 23% for the WP-condA category and 38% for the Ad-Pl. Unlike the drought model (model 2), model 3 assessing the impact of freeze conditioning on drought had much lower explanatory power and was not significant for any category.

**Table 1:** Redundancy analyses (RDA) based variance partitioning, model *R*^2^ and significance of the multivariate models along with representative candidate genes for the three *Q*_ST_ categories. Candidates were determined manually based on the annotations from EnTAP (see Table S1 & Methods S1) by focusing on transcripts with highest *Q_ST_* values in each category.

| Category | Model | $R^2$ | $R^2_{adj}$ | $p$-value | Candidates |
| --- | --- | --- | --- | --- | --- |
| BS-condA | Drought + Freeze + Geography<br>(model 1) | 0.88 | 0.5 | 0.003 | WRKY4, CYC4,<br>XTH9, LRR,<br>VIN3, CO-like,<br>PPR, ERF1,<br>GAST1, TIP1 |
|  | Drought + Geography<br>(model2) | 0.77 | 0.48 | 0.001 |  |
|  | Freeze + Geography<br>(model3) | 0.45 | 0.02 | 0.4 |  |
| WP-condA | Drought + Freeze + Geography<br>(model1) | 0.81 | 0.17 | 0.2 | CyP450, ARF2,<br>ERF2, SWEET, |
|  | Drought + Geography<br>(model2) | 0.65 | 0.23 | 0.04 | Ycf3, WRKY,<br>PPR, LRR |
|  | Freeze + Geography<br>(model3) | 0.5 | 0.1 | 0.17 |  |
| Adap-PI | Drought + Freeze + Geography<br>(model1) | 0.82 | 0.19 | 0.3 | SAUR50, α-<br>expansin, ERG3,<br>LPPD |
|  | Drought + Geography<br>(model2) | 0.68 | 0.38 | 0.03 |  |
|  | Freeze + Geography<br>(model3) | 0.34 | 0 | 0.8 |  |

Using 23,261 ddRADseq markers, estimates of genomic ancestry for the 30 maternal trees ranged from 0.18 to 1 (Fig. S2b), with a value of 1 indicating 100% genomic ancestry from *Pinus flexilis*. Proportion of sibs surviving per maternal family ranged from 0.37 to 1 in BS and from 0.23 to 0.90 in WP. Across both gardens, estimated survival of maternal family was positively correlated with genomic ancestry (Pearson’s *r* for WP = 0.24, p = 0.18 and for BS = 0.40, p=0.02). Across all transcripts, population level expression values were weakly correlated with population level estimates of ancestry (−0.02 ± 0.31 in BS and 0.02 ± 0.31 in WP). The distribution of correlation coefficients for the relationship between mean population genomic ancestry and population level expression values ranged from −0.6 to 0.8 for BS-condA, from −0.8 to 0.8 for WP-condA and from −0.6 to 0.4 for Ad-Pl. While the observed distribution was slightly shifted towards negative values for Ad-Pl, the cumulative distribution of correlation coefficients between ancestry and transcript abundance for all categories was not significantly different from their respective matched background set of transcripts (BS-condA p= 0.702 & D = 0.046, WP-condA p = 0.84 & D = 0.03, Ad-Pl p = 0.74 & D = 0.13).

Using a similar approach we evaluated the relationship between population level expression values and estimated maternal survival for each of the *Q_ST_* categories. Overall, expression values were weakly correlated with our estimate of survival (−0.008 ± 0.32 in BS & 0.029 ± 0.33 in WP). Correlation coefficients ranged from −0.68 to 0.76 for BS-condA, from - 0.89 to 0.73 for WP-condA and from −0.42 to 0.55 for Ad-Pl category. Similar to our observation for the relationship between ancestry and transcript abundance, the observed cumulative distribution of correlation coefficients for all categories was not significantly different from their respective backgrounds (BS-condA p= 0.817 & D = 0.073, WP-condA p = 0.11 & D = 0.065, Ad-Pl p= 0.39 & D = 0.17).

No functional groups, as assessed using GO terms, were significantly enriched across the three *Q_ST_* categories. In contrast, GO terms related to signal transduction and cell communication were significantly depleted for BS-condA, while GO terms related to hydrolase activity were significantly depleted for WP-condA (Table S2). By manually scanning the annotations for each category (Table S1), we note several transcripts related to freeze and drought tolerance as wekk as cell wall modulation (Sung & Amasino, 2005; Raimund, 2015; Tucker et al. 2018) such as *VIN3, PPR, XTH* and *Aquaporins* like TIP1 in the BS-condA category. Transcripts in the WP-condA category were notably related to drought stress response and photosynthesis (Lata et al. 2015; Zhang et al. 2020), such as several *ERF* family genes, *SWEET, WRKY* and *psbE*, as well as auxin responsive elements such as *ARF2*. Finally, for transcripts in the Ad-Pl category, we noted auxin responsive elements such as SAUR50 (Sun et al. 2016) critical for light signaling, α-expansin and the ABA signaling pathway family genes such as LPPD which have known roles in several plant developmental and stress response pathways (Marowa et al. 2016).

### Drivers of adaptive evolution at the co-expression module level (H2 & H3)

At the module level, across both gardens we noted a wide range of eigengene associations with ancestry, ranging from −0.56 to 0.35 in WP and −0.32 to 0.41 in BS (Table S3). Compared to the association with ancestry, eigengene expression was generally more strongly related to survival and ranged from −0.50 to 0.72 in both gardens (Table S3). In WP the module ME11 exhibited a significant (p < 0.05) negative association with ancestry, as well as with survival (Table 2), however the association with ancestry was not significant after using a false discovery rate (FDR) of 0.05. In BS, ME13 exhibited a significant (p < 0.05) positive association with ancestry and survival, but was noted in WP the association with ancestry was not significant after FDR correction (Table 2).

**Table 2:** Candidate multivariate traits or MEs associated with survival, ancestry or exhibiting *Q*_ST_ enrichment along with the annotation of the transcript with the highest or the next highest module membership (hMEM) in cases where the top most transcript was lacking a clear annotation.

| Garden | Module ID | Survival association | Ancestry association | $Q_{ST}$ enrichment value | hMEM annotation |
| --- | --- | --- | --- | --- | --- |
| BS | ME13 | 0.62<br>($p = 2e-04$,<br>$q = 0.003$) | 0.41<br>($p = 0.02$,<br>$q = 0.23$) | 2.87 | Golgi intermediate protein 3 |
|  | ME26 | 0.71<br>(p = 1e-05,<br>q = 3e-04) | 0.29<br>(p = 0.01,<br>q = 0.23) | 0.644 | WRKY14 |
|  | ME16 | -0.19<br>(p = 0.32) | -0.17<br>(p = 0.36) | 1.37 | Non-race specific<br>disease resistance-1<br>(NDR1) |
|  | ME12 | -0.18<br>(p = 0.34) | -0.11<br>(p = 0.55) | 2.12 | Snakin-2 |
|  | ME14 | 0.29<br>(p = 0.12) | 0.049<br>(p = 0.79) | 2.12 | E3 ubiquitin-protein<br>ligase (KEG) |
| WP | ME24 | 0.53<br>(p = 0.003,<br>q = 0.016) | 0.23<br>(p = 0.22) | 3.96 | β-carotene hydrolase<br>(CHY-β) |
|  | ME19 | 0.69<br>(p = 3e-05,<br>q = 0.0003) | 0.35<br>(p = 6e-02,<br>q = 0.40) | 0.99 | Light regulated WD1 |
|  | ME8 | -0.27<br>(p = 0.2) | -0.56<br>(p = 0.001,<br>q = 0.027) | 0 | Trehalose-6-phosphate<br>synthase 1 |
|  | ME11 | -0.43<br>(p = 0.02,<br>q = 0.04) | -0.47<br>(p = 0.009,<br>q = 0.12) | 1.62 | E3-ubiquitin protein<br>ligase (UPL5-like) |
|  | ME2 | 0.43<br>(p = 0.02,<br>q = 0.04) | 0.17<br>(p = 0.36) | 1.41 | Phospholipase C Gene<br>Family- like (PI-PLC) |
|  | ME1 | -0.24<br>(p = 0.2) | -0.017<br>(p = 0.9) | 2.10 | RNA binding protein |
|  | ME3 | -0.22<br>(p = 0.23) | 0.11<br>(p = 0.6) | 2.02 | Filament like protein<br>(FPP7) |
|  | ME15 | 0.3<br>(p = 0.08) | 0.21<br>(p = 0.3) | 1.72 | ubiquitin-like modifier<br>(ULP) |

GO enrichment analysis for modules exhibiting *Q*_ST_ category enrichment, depletion, strong association with ancestry or with survival identified a wide range of functional groups across both gardens (Table 2, Table S2). Nevertheless, some notable patterns emerged. Across both gardens, *Q*_ST_ category enrichment modules were predominantly associated with GO terms and genes related to stress response, cell wall modulation and response to light stimuli. In contrast, *Q*_ST_ category depleted modules across gardens were predominantly represented by developmental and metabolic processes. One notable exception to this pattern was the ME12 module in BS which exhibited a strong enrichment but was associated with floral and reproductive processes. In WP the ME16 module was associated with similar functional terms as ME12 in BS but exhibited a strong depletion for our garden specific *Q*_ST_ category. Functional terms associated with metabolic processes were generally under-represented, and terms associated with response to abiotic stress were over-represented in the modules strongly associated with both survival and ancestry in WP (ME11) as well as in BS (ME13). Across both gardens, modules exhibiting the strongest association with survival (ME26 in BS and ME19 in WP) were primarily associated with GO terms related to glutathione transferase, detoxification and response to abiotic stimuli.

### Relationship between network connectivity and adaptive evolution (H4)

Across both gardens, *Q_ST_* was positively associated with module connectivity (kWithin Pearson’s *r* BS = 0.36, p < 0.0001 & WP = 0.46, p < 0.0001) and with total network connectivity (kTotal Pearson’s *r* BS =0.53, p < 0.0001& WP = 0.56, p < 0.0001). Additionally, both WP-condA and BS-condA transcripts as a group were significantly over-represented among transcripts located in the core of the module (obs WP = 1.74, p < 0.0001; obs BS = 2.5, p < 0.0001) as well at the core of the garden specific co-expression network (obs WP = 2.0, p < 0.0001; obs BS =3.9, p < 0.0001) (Fig. 4; Fig. S3). This pattern persisted even after accounting for differences in expression levels between the core and the periphery transcripts. For most modules exhibiting strong population differentiation, the transcript with highest *Q_ST_* also had the highest module membership and hence was located at the module core (Fig. S3). However, the average *Q_ST_* for core transcripts was higher than that of the periphery only for 38% of the strong differentiated modules in BS and for 20% of the differentiated modules in the WP (Table S3). Most modules were preserved across gardens with respect to either *medianRank* or *Zsummary*. Our stringent threshold using the union of both measures identified five modules out of the 31 identified in BS as being strongly preserved in WP and four out of the 26 WP modules being strongly preserved in BS (Table S3; Fig. 4). Although overall co-expression network preservation was high across gardens, some interesting module specific patterns emerged. Modules enriched for *Q_ST_* transcripts usually had higher *medianRank* and were less preserved across gardens while modules depleted for *Q_ST_* had consistently lower *medianRank* and often strongly preserved across gardens (Fig. 4a,c).

## Discussion

### Hybrid zone populations exhibit adaptive potential to novel climatic conditions

Long-lived tree species in the northern hemisphere have experienced long-term climatic fluctuations causing them to be distant from their current climate optima. As a result, the present patterns of genetic diversity are often structured by past environmental events (Hampe & Petit, 2005; Mayol et al. 2015) causing populations to exhibit an adaptation lag and not necessarily be devoid of potential adaptive variants. Theory suggests that GEI should be prevalent in populations exhibiting an adaptation lag and occurring in heterogenous environments (Via & Lande, 1985; Ghalambor et al. 2007). Sequential founder events, high levels of landscape fragmentation and the associated loss of genetic diversity, however, can restrict the evolution of GEI (Schmid et al. 2019).

The populations used in this study occur on fragmented landscapes in the sky islands of the southwestern United States, where they experience marked seasonal and annual fluctuations in environmental conditions as well as gene flow from a northern sister species, *P. flexilis* (Menon et al. 2018). By conceptualising gene expression and co-expression modules formed by them as a quantitative trait we revealed strong signals of GEI and lend support to H1 (Fig. 2, 4). However, in relation to studies conducted in conifers (reviewed in Lind et al. 2018) we found a larger number of strongly differentiated traits (Fig. 3). This could be a result of sampling the transcriptome profile of juvenile trees representing a hybrid zone. Selective pressures, specifically those involved in post-zygotic isolating barriers, tend to be strongest during early life stages across tree species (Lindtke et al. 2014; Zhao et al. 2014). Additionally, trait differentiation will be elevated during early stages of species divergence (Ogasawara & Okubo, 2009). Taken together this suggests that one possibility for the strongly differentiated transcripts identified here is their role in the maintenance of species boundaries between *P. flexilis* and *P. strobiformis*. Another possibility is our limited population sampling which resulted in high levels of uncertainty around point estimates of *Q_ST_* resulting in only 2.5% of transcripts being classified under the *Q_ST_* categories (Fig. 2) regardless of the U-shaped distribution we note for the point estimates (Fig. 3). The observed proportion of putatively adaptive traits identified through our permutation approach, however, agrees with estimates across other systems (Roberge et al. 2007; Leder et al. 2015). At the co-expression module level, we noticed similarly strong signals of population differentiation (Table S3). While some of these modules could be false positives due to population structure in the gene expression dataset used to construct co-expression modules, eight of the strongly differentiated modules were also enriched for our target *Q_ST_* categories (Fig. 4). We suggest these are likely true candidates for garden-specific signatures of adaptive evolution, specifically given the overall weak population structure in our study system (Menon et al. 2018).

Taken together the strong GEI noted in our study likely resulted from a combination of juvenile trees exhibiting environment-dependent hybrid fitness, high heterogeneity and strong differences in source environmental conditions (Hopken et al. 2013; Williams et al. 2010) and the novel selective environment the seedlings were exposed to. The signals of local adaptation noted under novel environmental conditions in our study design suggest that long-lived species such as conifers may not be limited in their ability to adapt rapidly to changing climatic conditions. At the very least, these populations have ample standing variation despite their fragmented habitat that could aid response to novel selective pressures.

### Drought and hybrid ancestry influence adaptive evolution and generate GEI

Results from our transcript categorisation and co-expression module enrichment analyses partially agreed with H2. While we were able to identify garden specific patterns of adaptive differentiation and some level of sharing at both the per-transcript (Fig. 2) and co-expression module level (Fig. 4), the driver of adaptive evolution remained the same across gardens. Even though BS is much colder than WP, both the WP-condA and BS-condA sets were significantly associated only with drought and not freeze-related variables (Table 1). Similarly, at the co-expression network level we noted high overall preservation across gardens, despite the *Q*_ST_ enriched modules exhibiting weak preservation as would be expected given their garden specific categorisation. There are several possible explanations for this. First, the high preservation could be reflective of physiological and molecular overlap and interactions between mechanisms associated with drought and cold tolerance (Pandey et al. 2015; Zhou et al. 2019). Second, trees across both gardens experienced a 50% reduction in ambient precipitation even though generally BS was colder and less dry than WP. This shared drought stress implemented both as part of our space-time-substitution design and artificially through rain-out shelters could have generated the high network preservation and a strong impact of drought on adaptive trait differentiation. Third, in agreement with previous work demonstrating drought as a major selective gradient in these populations (Goodrich et al. 2016; Bucholz et al. 2020; Menon et al. 2021) and strong genetic control for growth cessation phenology in trees (Howe et al. 2007; Holliday et al. 2010; Swenson et al. *in prep*), our environment association results reflect a stronger impact of source population selective environment on trait differentiation (Table 1). Specifically, had we sampled during bud flush and the active growth period following it, which is known to be under lower genetic control, we might have noted lower network preservation and perhaps a marked impact of freeze stress. Finally, network preservation statistics are predominately assessed using patterns of connectivity among transcripts, causing the noted preservation to be likely driven by the importance of pleiotropic architecture across both gardens that exposed the trees to extreme environmental conditions (Fig. 1.b).

At the multivariate level, gene expression modules associated with ancestry were often also strongly associated with survival (Table S3; Table 2). Specifically, two *Q_ST_* enriched modules - ME24 in WP and ME13 in BS-were strongly associated with *P. flexilis* ancestry as well as with survival making the genes and regulatory regions within these modules a key candidate for further detailed studies of GEI (Fig. 4; Table 2). The absence of strong associations with ancestry for the three *Q_ST_* categories is likely because our study utilised estimates of ancestry from ddRAD-seq dataset, which although is representative of overall genomic ancestry, provides less coverage of genic regions in species with large genomes such as pines (Parchman et al. 2018). Interestingly, *P. flexilis* ancestry was positively associated with survival across both gardens. Despite this positive trend, the association between survival and ancestry was significant only in BS and both survival as well as ancestry remained uncorrelated to population transcript abundance levels. These results thus provide only partial support for H3 and highlight the need for finer scale evaluation to gather more evidence for hybrid ancestry driving GEI. However, drawing from our previous work (Menon et al. 2021) and a likely wider environmental tolerance of *P. flexilis* than *P. strobiformis* (Windmuller-Campione & Long, 2016) we speculate that significant association between ancestry and survival only in BS is indicative of *P. flexilis* like variants enhancing survival in cooler environments. Overall, by using a space-for-time substitution study design and measuring several quantitative traits through transcriptomics, we highlight that conifer hybrid zones contain substantial additive genetic variation for expression traits with a likely environment dependent contribution of introgressed variants.

### Transcriptome wide profiling illustrates the prevalence of pleiotropic architecture

Strongly connected core genes within biological networks often experience selective constraints, as mutations occurring in these genes impart pleiotropic effects which could manifest as deleterious to multiple downstream traits (Jordan et al. 2004). On the other hand, genes with lower connectivity will often be involved in adaptive evolution and in GEI as selection can fine tune population responses to local environment without disrupting preserved functional components (Cork & Purugganan, 2004; Josephs et al. 2017). Despite this, our work demonstrates an overrepresentation of traits exhibiting GEI in the core of the network and at the core of the module, thereby rejecting H4. Following Fisher’s geometric model (Fisher 1930), if populations are further from their fitness optima, large effect loci that impart pleiotropic influences might be advantageous during the initial stages of the adaptive walk (Orr 1998). We suggest that this is likely the case for our expanding hybrid zone populations (Menon et al. 2020) that were exposed to novel selective regimes. While further validation through eQTL analysis is warranted, similar enrichment of strongly differentiated transcripts in the core of the co-expression network has been noted in other studies (des Marias et al. 2017; Hamala et al. 2020; Chateigner et al. 2020) and hence is unlikely an anomaly.

## Conclusions

Species responses to changing climatic conditions are variable across populations and involve a combination of adaptation, migration and extirpation (Aitken et al. 2008). Common garden approaches and genome-wide association studies across plants have provided remarkable evidence for GEI and for local adaptation (Langlet, 1971; Savolainen et al. 2013). Our results show strong signals of GEI at both per-transcript and co-expression module level reiterating decades of studies demonstrating local adaptation in forest trees. By utilizing a space-for-time substitution design we additionally demonstrate that long-lived tree species are likely equipped to respond to rapidly changing climatic conditions through the initial advantage provided by large effect pleiotropic loci. Additionally, we highlight the environment dependent impact of interspecific gene flow towards adaptive evolution in the *P. strobiformis-P. flexilis* hybrid zone populations. Our work stands among recent efforts to evaluate adaptive potential of species at or beyond their climate edge (Geber & Eckhart, 2005; Olsen et al. 2019) and highlights the utility of transcriptomics to assess the architecture aiding evolution towards a novel climate optimum.

## Supporting information

**Methods S1:** Details of the pipeline and summaries used for denovo transcriptome assembly.

**Methods S2:** Pipeline details for SNP calling and filtering using ddRAD-seq genotyping.

**Methods S3**: Details of steps and summary statistics for construction of the co-expression networks.

**Figure S1: a)** PCA of 76 environmental variables from climateWNA for the normal 1981-2010 for the 10 sampled populations and for the study year 2019 for the common gardens. **(b)** Heatmap representing the proportion of times drought or freeze related environmental variables measured at the gardens in 2019 were outside the range experienced by the assayed population across a 60 year period. Garden environment was declared as an outlier if it fell outside the 95 percentage confidence interval of the climatic condition experienced by the populations for each year. This evaluation was done on a year-by-year basis. Warmer (red) colours indicate higher drought intensity and colder (blue) colours indicate higher freeze events experienced by the gardens relative to the assayed populations.

Variables used for **(b)** Tmax_wt = winter mean max temperature, Tmin_wt = winter mean min temperature, Tave_wt = winter mean temperature, DD_0_wt = winter degree-days below 0°C, DD5_wt = winter degree-days below 5°C, DD_18_wt= winter degree-days below 18°C, DD18_wt=winter degree-days above 18°C, NFFD_wt=winter frost-free days, PAS_wt=winter precipitation as snow (mm), Eref_wt = winter Hargreaves reference evaporation, Tmax_sm = summer mean max temperature, Tmin_sm = summer mean min temperature, Tave_sm = summer mean temperature, DD5_sm = summer degree-days above 5°C, DD18_sm = summer degree-days above 18°C, DD_18_sm= summer degree-days below 18°C, Eref_sm= summer Hargreaves reference evaporation (mm), CMD_sm = summer Hargreaves climatic moisture deficit,RH_sm= summer relative humidity (%), CMI_sm = summer Hogg’s climate moisture index (mm), PPT_sm = summer precipitation.

**Figure S2: a)** Geographical range map of *P. strobiformis* and *P. flexilis* along with the populations sampled for ddRAD-seq genotyping. **b)** STRUCTURE plot of genomic ancestry for the 30 maternal trees represented by the mixed-sibs sampled for RNA-seq and **c)** Distribution of F_ST_ values for the 30 maternal trees.

**Fig S3:** Visualization of *Q*_ST_ enriched modules in BS and WP with circles representing the nodes (transcripts) and lines representing the edges (correlation between transcripts). Transcripts classified as BS-condA or WP-condA are highlighted in color while the rest are represented by black circles. The location of the transcripts in the module are representative of their edge weights (i.e intra-modular connectivity).

**Fig S4:** Relationship between mean transcript expression and co-expression connectivity levels for **a)** WP and **b)** BS. Transcripts were divided into bins of mean expression by their position in the co-expression network (core vs. periphery) to highlight their level of connectivity.

**Figure S5: A)** Evaluation of scale free topology at various soft-thresholding powers and **B)** clustering dendrogram of transcripts using a soft-thresholding of 8 for BS and 9 for WP.

**Table S1**: Annotations for all 47,651 transcripts generated through EnTAP.

**Table S2:** Summary of GO terms and the enrichment analyses for three *Q*_ST_ categories (BS-condA, WP-condA, Ad-Pl) and for all modules across the two gardens.

**Table S3:** Summary statistics for modules identified across the gardens. Columns represent eigengene association with survival and ancestry, *Q*_ST_ enrichment score, annotation of the transcript with the highest module membership, preservation scores and difference in mean *Q*ST between transcripts classified in the core and periphery of a module (*Q_ST_* periphery - *Q_ST_* core).

## REFERENCES

1. Adams HD, Kolb TE. 2004. Drought responses of conifers in ecotone forests of northern Arizona: tree ring growth and leaf δ 13 C. Oecologia. 140: 217–225.

2. Aitken SA, Yeaman S, Holliday JA, Wang T, Curtis-McLane S. 2008. Adaptation, migration or extirpation: climate change outcomes for tree populations. Evolutionary Applications. 1: 95–111.

3. Bates D, Mächler M, Bolker B, Walker S. 2015. Fitting Linear Mixed-Effects Models Using lme4. Journal of Statistical Software, 67(1), 1–48.

4. Bisbee J. 2014. Cone morphology of the Pinus ayacahuite-flexilis complex of the southwestern United States and Mexico. Bulletin of the Cupressus conservation project, 3, 3–33.

5. Bradshaw AD. 2006. Unravelling phenotypic plasticity–why should we bother? New Phytologist 170: 644–64.

6. Brooks ME et al. 2017. Modeling zero-inflated count data with glmmTMB. BiorXiv. https://doi.org/10.1101/132753.

7. Bucholz ER, Waring KM, Kolb TE, Swenson JK, and Whipple AV. 2020. Water relations and drought response of *Pinus strobiformis*. Canadian Journal of Forest Research. 50(9): 905–916.

8. Chateigner A., Lesage-Descauses MC., Rogier O. et al. 2020. Gene expression predictions and networks in natural populations supports the omnigenic theory. BMC Genomics 21: 416 (2020).

9. Clausen Jens, Keck David D, Hiesey William M. 1948. Experimental studies on the nature of species. III: Environmental responses of climatic races of Achillea. 581; Washington, D.C.: Carnegie Institution of Washington.

10. Cook B, Ault T and Smerdon JE. 2015. Unprecedented 21st century drought risk in the American Southwest and Central Plains. Science Advances. 1:e1400082

11. Cork JM, Purugganan MD. 2004. The evolution of molecular genetic pathways and networks. Bioessays. 26(5):479–84.

12. Danecek P, et al. 2011. The Variant Call Format and VCFtools, Bioinformatics.

13. Dayan DI, Crawford DL, Oleksiak MF. 2015. Phenotypic plasticity in gene expression contributes to divergence of locally adapted populations of Fundulus heteroclitus. Molecular Ecology. 24(13):3345–59.

14. Des Marais DL, Guerrero RF, Lasky JR, Scarpino SV. 2017. Topological features of a gene co-expression network predict patterns of natural diversity in environmental response. Proceedings of Royal Society of Biological Sciences. 284(1856):20170914

15. Falconer DS & Mackay T. 1996. Introduction to Quantitative Genetics. Longman, New York.

16. Fisher RA. 1930. The genetic theory of natural selection. Oxford: The Carendon Press.

17. Geber MA, Eckhart VM. 2005. Experimental studies of adaptation in Clarkia xantiana. II. Fitness variation across a subspecies border. Evolution 59(3):521–31.

18. Ghalambor CK, McKay JK, Carroll SP, Reznick DN. 2007. Adaptive versus non-adaptive phenotypic plasticity and the potential for contemporary adaptation in new environments. Functional ecology. 2007. 21(3):394–407.

19. Goodrich BA, Waring KM & Kolb TE. 2016. Genetic variation in *Pinus strobiformis* growth and drought tolerance from southwestern US populations. Tree Physiology. 36: 1219–1235.

20. Goudet J. 2005. hierfstat, a package for R to compute and test hierarchical F-statistics. Molecular Ecology Notes, 5, 184–186.

21. Grabherr MG et al. 2013. Trinity: reconstructing a full-length transcriptome without a genome from RNA-Seq data. Nature Biotechnology. 29(7): 644–652.

22. Grote S. 2020. GOfuncR: Gene ontology enrichment using FUNC. R package version 1.10.0.

23. Haas BJ et al. 2014. *De novo* transcript sequence reconstruction from RNA-Seq: reference generation and analysis with Trinity. Nature Protocols. 8: 1–43.

24. Hämälä T, Guiltinan MJ, Marden JH, Maximova SN, dePamphilis CW & Tiffin P. 2020.a. Gene Expression Modularity Reveals Footprints of Polygenic Adaptation in *Theobroma cacao*. Molecular Biology and Evolution. 27(1):110–123.

25. Hämälä T, Gorton AJ, Moeller DA, Tiffin P. 2020.b. Pleiotropy facilitates local adaptation to distant optima in common ragweed *(Ambrosia artemisiifolia)*. PLOS Genetics 16(3): e1008707.

26. Hampe A. & Petit RJ. 2005. Conserving biodiversity under climate change: the rear edge matters. Ecology Letters. 8:461–467.

27. Hart AJ et al. 2020. EnTAP: Bringing faster and smarter functional annotation to non-model eukaryotic transcriptomes. Molecular Ecology Resources. 20: 591–604.

28. Hartwell LH, Hopfield JJ, Leibler S, Murray AW (1999). From molecular to modular cell biology. Nature. 402: C47–52.

29. Hoffman GE and Schadt EE. 2016. variancePartition: interpreting drivers of variation in complex gene expression studies. BMC Bioinformatics 17: 483.

30. Holliday JA, Ritland K, & Aitken SN. 2010. Widespread, ecologically relevant genetic markers developed from association mapping of climate-related traits in Sitka spruce (*Picea sitchensis*). The New Phytologist. 188, 501–514.

31. Hopken MW, Douglas MR, Douglas ME. 2013. Stream hierarchy defines riverscape genetics of a North American desert fish. Molecular Ecology. 22: 956–971.

32. Howe GT, Aitken SN, Neale DB et al. 2003. From genotype to phenotype: unraveling the complexities of cold adaptation in forest trees. Canadian Journal of Botany 81: 1247–1266.

33. Huang W, Carbone MA, Lyman RF. et al. 2020. Genotype by environment interaction for gene expression in *Drosophila melanogaster*. Nature Communications. 11: 5451.

34. Jordan IK, Mariño-Ramírez L, Wolf YI & Koonin EV. 2004. Conservation and Coevolution in the Scale-Free Human Gene Coexpression Network, Molecular Biology and Evolution, 21(11):2058–2070.

35. Josephs EB, Wright SI, Stinchcombe JR, Schoen DJ. 2017. The relationship between selection, network connectivity, and regulatory variation within a population of Capsella grandiflora. Genome Biology Evolution. 9(4):1099–1109.

36. Krueger F. Trim Galore!: 2015. A wrapper tool around Cutadapt and FastQC to consistently apply quality and adapter trimming to FastQ files. http://www.bioinformatics.babraham.ac.uk/projects/trim_galore/

37. Langlet O. 1971. Two hundred years of genecology. Taxon 2:653–721

38. Langfelder P, Horvath S. 2008. WGCNA: an R package for weighted correlation network analysis. BMC Bioinformatics. 9:559.

39. Langfelder P, Luo R, Oldham MC, Horvath S. 2011. Is My Network Module Preserved and Reproducible? PLoS Computational Biology. 7(1): e1001057.

40. Langmead B & Salzberg S. 2012. Fast gapped-read alignment with Bowtie 2. Nature Methods. 9:357–359.

41. Lasky JR, Des Marais DL, Lowry DB et al. 2014. Natural Variation in Abiotic Stress Responsive Gene Expression and Local Adaptation to Climate in Arabidopsis thaliana. Molecular biology and evolution. 31: 2283–2296.

42. Lata C., Muthamilarasan M., Prasad M. 2015. Drought Stress Responses and Signal Transduction in Plants. In: Pandey G. (eds) Elucidation of Abiotic Stress Signaling in Plants. Springer, New York, NY.

43. Law CW, Chen Y, Shi W. et al. 2014. voom: precision weights unlock linear model analysis tools for RNA-seq read counts. Genome Biology. 15: R29.

44. Leder EH, McCairns RJ, Leinonen T, Cano JM, Viitaniemi HM, Nikinmaa M, Primmer CR, Merilä J. 2015. The Evolution and Adaptive Potential of Transcriptional Variation in Sticklebacks—Signatures of Selection and Widespread Heritability. Molecular Biology and Evolution. 32(3):674–689.

45. Leinonen T, O’Hara RB, Cano JM and Merilä J. 2008. Comparative studies of quantitative trait and neutral marker divergence: a meta-analysis. Journal of Evolutionary Biology. 21:1–17.

46. Lind BM, Menon M, Bolte CE, Faske TM & Eckert AJ. 2018. The genomics of local adaptation in trees: Are we out of the woods yet?. Tree genetics & genomes. 14(2): 29.

47. Lindtke D, Gompert Z, Lexer C, Buerkle CA. 2014. Unexpected ancestry of Populus seedlings from a hybrid zone implies a large role of postzygotic selection in the maintenance of species. Molecular Ecology. 23: 4316–4330.

48. Mähler N, Wang J, Terebieniec BK, Ingvarsson PK, Street NR, Hvidsten TR. 2017. Gene co-expression network connectivity is an important determinant of selective constraint. PLoS Genetics. 13(4):e1006402.

49. Marowa P, Ding A, Kong Y. 2016. Expansins: roles in plant growth and potential applications in crop improvement. Plant Cell Reports. 5(5):949–965.

50. Mayol M, Riba M, Gonzalez-Martinez SC et al. 2015. Adapting through glacial cycles: insights from a long-lived tree (Taxus baccata). New Phytologist. 208: 973–986.

51. Menon M, Landguth E, Leal-Saenz A et al. 2019. Tracing the footprints of a moving hybrid zone under a demographic history of speciation with gene flow. Evolutionary Applications. 13(1): 195–209.

52. Menon M, Bagley JC, Page GFM. et al. 2021. Adaptive evolution in a conifer hybrid zone is driven by a mosaic of recently introgressed and background genetic variants. Communications Biology. 4:160

53. Menon M et al. 2018. The role of hybridization during ecological divergence of southwestern white pine *(Pinus strobiformis)* and limber pine *(P. flexilis)*. Molecular Ecology. 27: 1245–1260.

54. Nicotra AB, Atkin OK et al. 2010. Plant phenotypic plasticity in a changing climate. Trends in Plant Science. 15(12):684–692 (2010).

55. Pandey P, Ramegowda V, Senthil-Kumar M. 2015. Shared and unique responses of plants to multiple individual stresses and stress combinations: physiological and molecular mechanisms. Frontiers in Plant Science. 16(6):723.

56. Pickles, B.J., Twieg, B.D., O’Neill, G.A., Mohn, W.W. and Simard, S.W., 2015. Local adaptation in migrated interior Douglas-fir seedlings is mediated by ectomycorrhizas and other soil factors. New Phytologist, 207(3): 858–871.

57. Pickett STA. 1989. Space-for-time substitution as an alternative to long-term studies. In: Long-term studies in ecology: approaches and alternatives. Springer-Verlag, New York, New York, USA. 110–135.

58. Roberge C, Guderley H and L. Bernatchez. 2007. Genome wide Identification of Genes Under Directional Selection: Gene Transcription QST Scan in Diverging Atlantic Salmon Subpopulations. Genetics 177: 1011–1022.

59. Savolainen O, Lascoux M, Merila J. 2013. Ecological genomics of local adaptation. Nature Reviews Genetics. 14: 807–820.

60. Schmid M, Dallo R & Guillaume F. 2019. Species’ range dynamics affect the evolution of spatial variation in plasticity under environmental change. The American Naturalist. 193: 798–813.

61. Skotte L Korneliussen TS & Albrechtsen A. 2013. Estimating individual admixture proportions from next generation sequencing data. Genetics 195: 693–702.

62. Spitze, K. 1993. Population-structure in Daphnia-obtusa - quantitative genetic and allozymic variation. Genetics 135: 367–374.

63. Sung S. & Amasino R.M. 2004. Vernalization in Arabidopsis thaliana is mediated by the PHD finger protein VIN3. Nature 427: 159–163.

64. Squillace AE. Average genetic correlations among offspring from open-pollinated forest trees. Silvae Genetica. 23: 149–156 (1974).

65. Swenson, JK. Menon M, Eckert AJ. (in prep). Detection of environment specific alleles for phenology and survival at the edge of a conifer’s climate niche.

66. Tenhaken R. 2015 Cell wall remodeling under abiotic stress. Frontiers in Plant Science. 5:771.

67. Tohge T, Nishiyama Y, Hirai MY et al. 2005. Functional genomics by integrated analysis of metabolome and transcriptome of Arabidopsis plants over-expressing an MYB transcription factor. Plant Journal. 42(2):218–35.

68. Tucker MR, Lou H, Aubert MK, et al. 2018. Exploring the Role of Cell Wall-Related Genes and Polysaccharides during Plant Development. Plants (Basel). 7(2):42.

69. O’Brien AM, Sawers RJH, Strauss SY, Ross-Ibarra J. 2019. Adaptive phenotypic divergence in an annual grass differs across biotic contexts. Evolution. 73(11):2230–2246.

70. Ogasawara O & Okubo K. 2009. On theoretical models of gene expression evolution with random genetic drift and natural selection. PLoS One 4:e7943

71. Oksanen J et al. 2013. vegan: Community Ecology Package. R package version 2.5-2.

72. Olsen J, Singh Gill G, Haugen R, Matzner SL, Alsdurf J, Siemens DH. 2019. Evolutionary constraint on low elevation range expansion: defense-abiotic stress-tolerance trade-off in crosses of the ecological model Boechera stricta. Ecology and Evolution. 9(20):11532–44.

73. Orr HA. 1998. The population genetics of adaptation: The distribution of factors fixed during adaptive evolution. Evolution. 52: 935–949.

74. Puritz JB, Hollenbeck CM & Gold JR. 2014. dDocent: a RADseq, variant-calling pipeline designed for population genomics of non-model organisms. PeerJ, 2: e431.

75. R Core Team. R v.3.6.3: A Language and Environment for Statistical Computing. Vienna, Austria: The R Foundation for Statistical Computing. R Foundation for Statistical Computing, Vienna, Austria (2020)

76. Robinson MD, McCarthy DJ, Smyth GK. 2010. edgeR: a Bioconductor package for differential expression analysis of digital gene expression data. Bioinformatics, 26(1), 139–140.

77. Via S, Lande R. 1985. Genotype-environment Interaction and The Evolution Of Phenotypic Plasticity. Evolution. 39(3):505–522.

78. Wang T, Hamann A, Spittlehouse DL, & Carroll C. 2016. Locally downscaled and spatially customizable climate data for historical and future periods for North America. PLoS One 11: e0156720.

79. Whitehead A and DL Crawford. 2006. Variation within and among species in gene expression: raw material for evolution. Molecular Ecology. 15: 1197–1211.

80. Williams PA, Allen CD, Millar CI, Swetnam TW, Michaelsen J, Still CJ, and Leavitt SW. 2010. Forest responses to increasing aridity and warmth in the southwestern United States. Proceedings of the National Academy of Sciences of the United States of America. 107(50):21289–21294.

81. Windmuller-Campione MA & Long JN, 2016. Limber Pine (Pinus flexilis James), a Flexible Generalist of Forest Communities in the Intermountain West. PloS One. 11(8), 1–16.

82. Wong, C.E., Li, Y., Whitty, B.R., Diaz-Camino, C., Akhter, S.R., Brandle, J.E., Golding, G.B., Weretilnyk, E.A., Moffatt, B.A. and Griffith, M., 2005. Expressed sequence tags from the Yukon ecotype of Thellungiella reveal that gene expression in response to cold, drought and salinity shows little overlap. Plant molecular biology, 58(4): 561–574.

83. Zhang Y, Zhang A, Li X, Lu C. 2020. The Role of Chloroplast Gene Expression in Plant Responses to Environmental Stress. International Journal of Molecular Sciences. 21(17):6082.

84. Zhao W, Meng J, Wang B et al. 2014. Weak Crossability Barrier but Strong Juvenile Selection Supports Ecological Speciation of the Hybrid Pine Pinus Densata on the Tibetan Plateau. Evolution. 68(11): 3120–3133.

85. Zhou R., Yu X., Zhao T. et al. 2019. Physiological analysis and transcriptome sequencing reveal the effects of combined cold and drought on tomato leaf. BMC Plant Biology. 19: 377

