## Supplemental figure 1 for "Variation in gene expression patterns across a conifer hybrid zone highlights the architecture of adaptive evolution under novel selective pressures"

A)

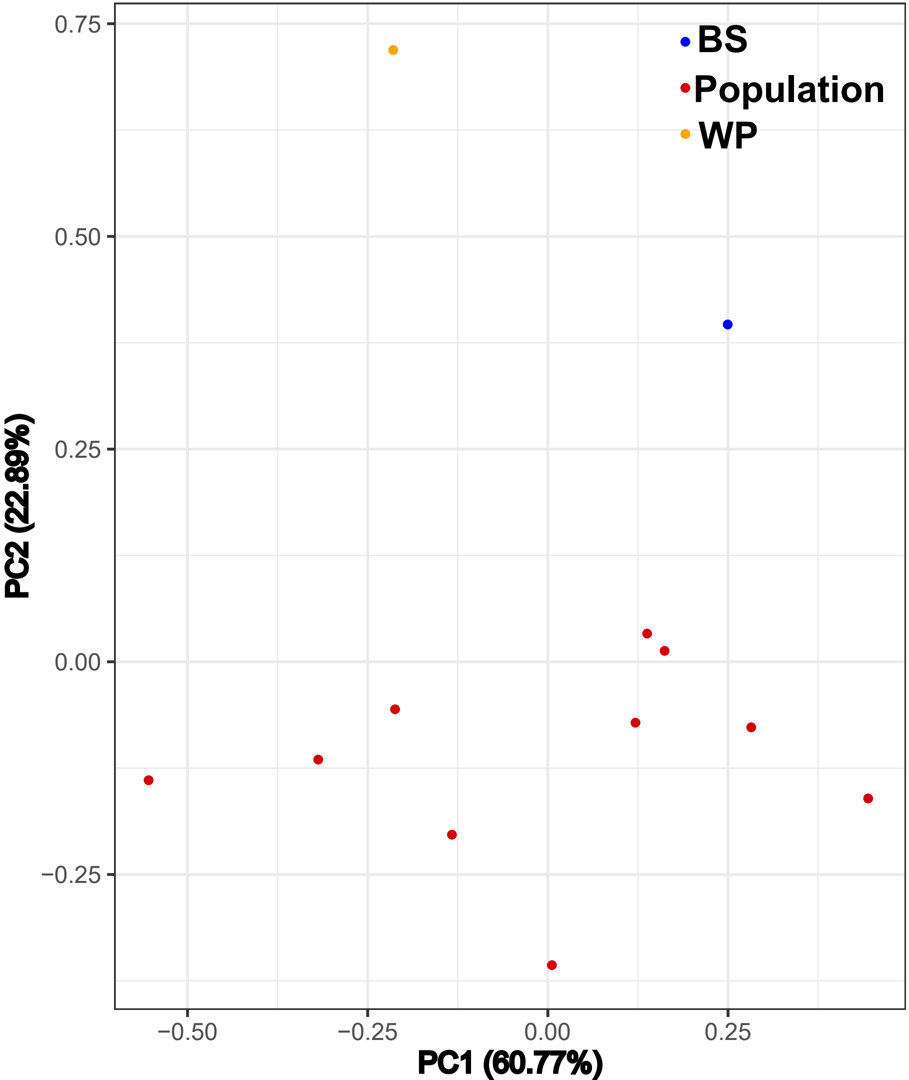


B)

| variable | BS | WP |
| --- | --- | --- |
| Tmax_wt | 100 | 0 |
| Tmin_wt | 0 | 0 |
| Tave_wt | 100 | 0 |
| DD_0_wt | 0 | 0 |
| DD5_wt | 63 | 0 |
| DD_18_wt | 0 | 0 |
| DD18_wt | 100 | 0 |
| NFFD_wt | 0 | 0 |
| PAS_wt | 100 | 0 |
| Eref_wt | 0 | 0 |
| Tmax_sm | 36 | 100 |
| Tmin_sm | 0 | 36 |
| Tave_sm | 0 | 100 |
| DD5_sm | 0 | 100 |
| DD18_sm | 0 | 100 |
| DD_18_sm | 0 | 100 |
| Eref_sm | 0 | 100 |
| CMD_sm | 100 | 100 |
| RH_sm | 0 | 0 |
| CMI_sm | 100 | 100 |
| PPT_sm | 100 | 100 |
