## Supplementary figures and images for "Variation in gene expression patterns across a conifer hybrid zone highlights the architecture of adaptive evolution under novel selective pressures"

### Supplemental figure 2

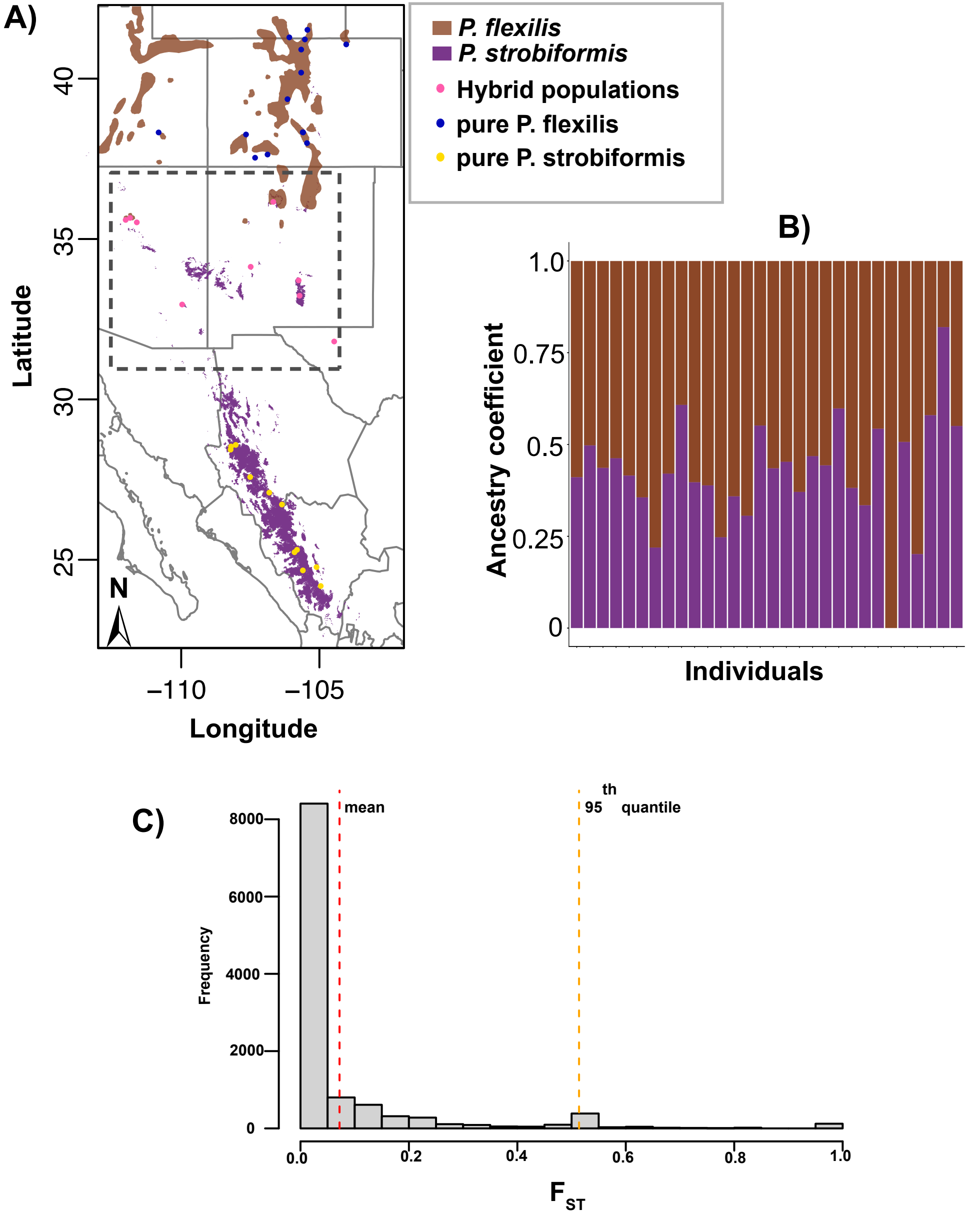

### Supplemental figure 3

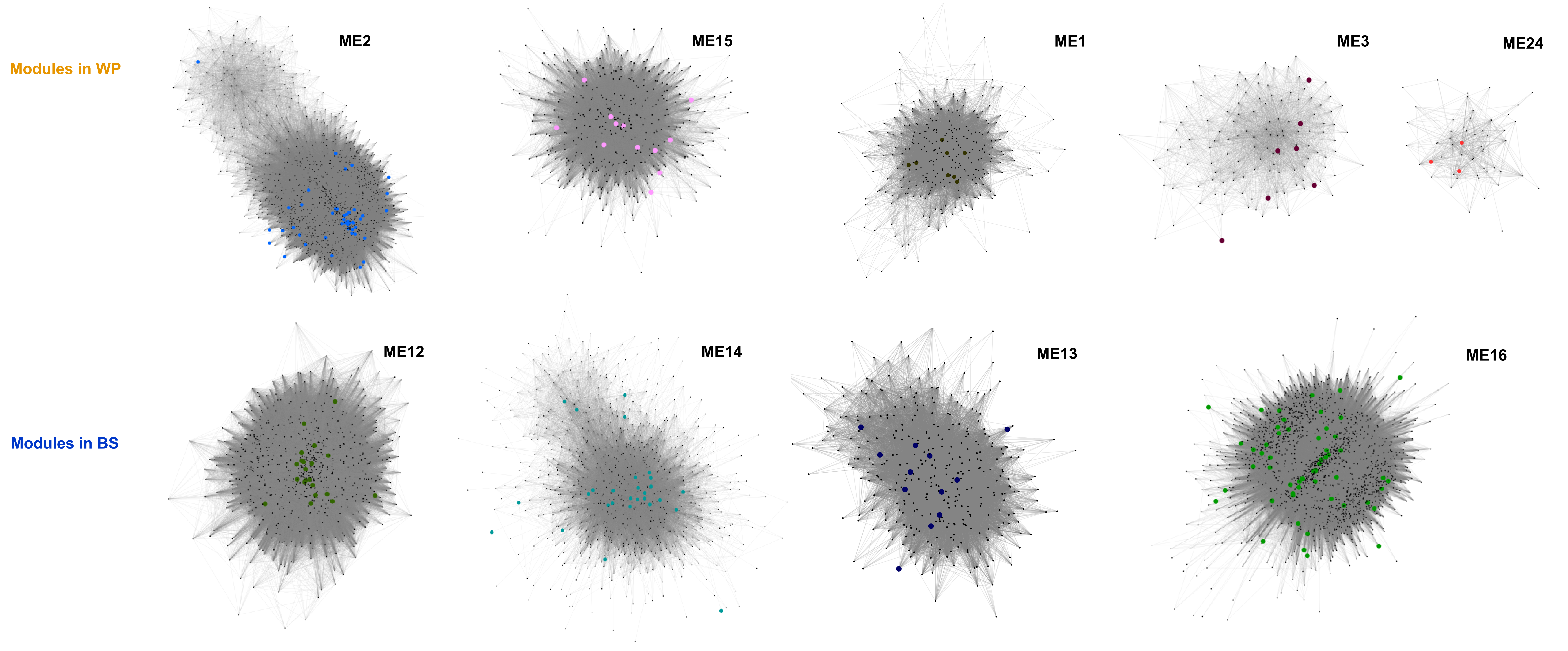

### Supplemental figure 4

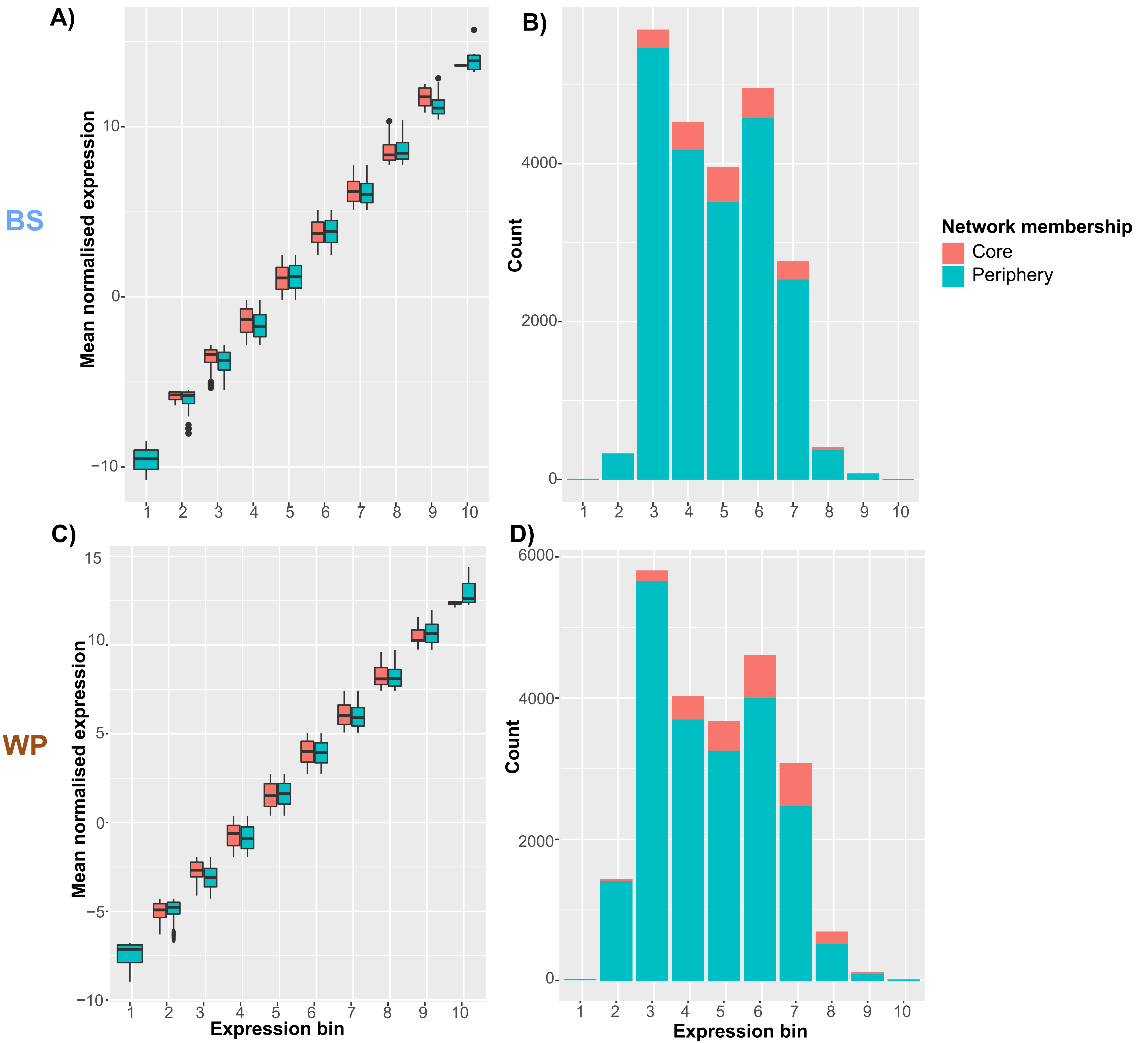

### Supplemental figure 5

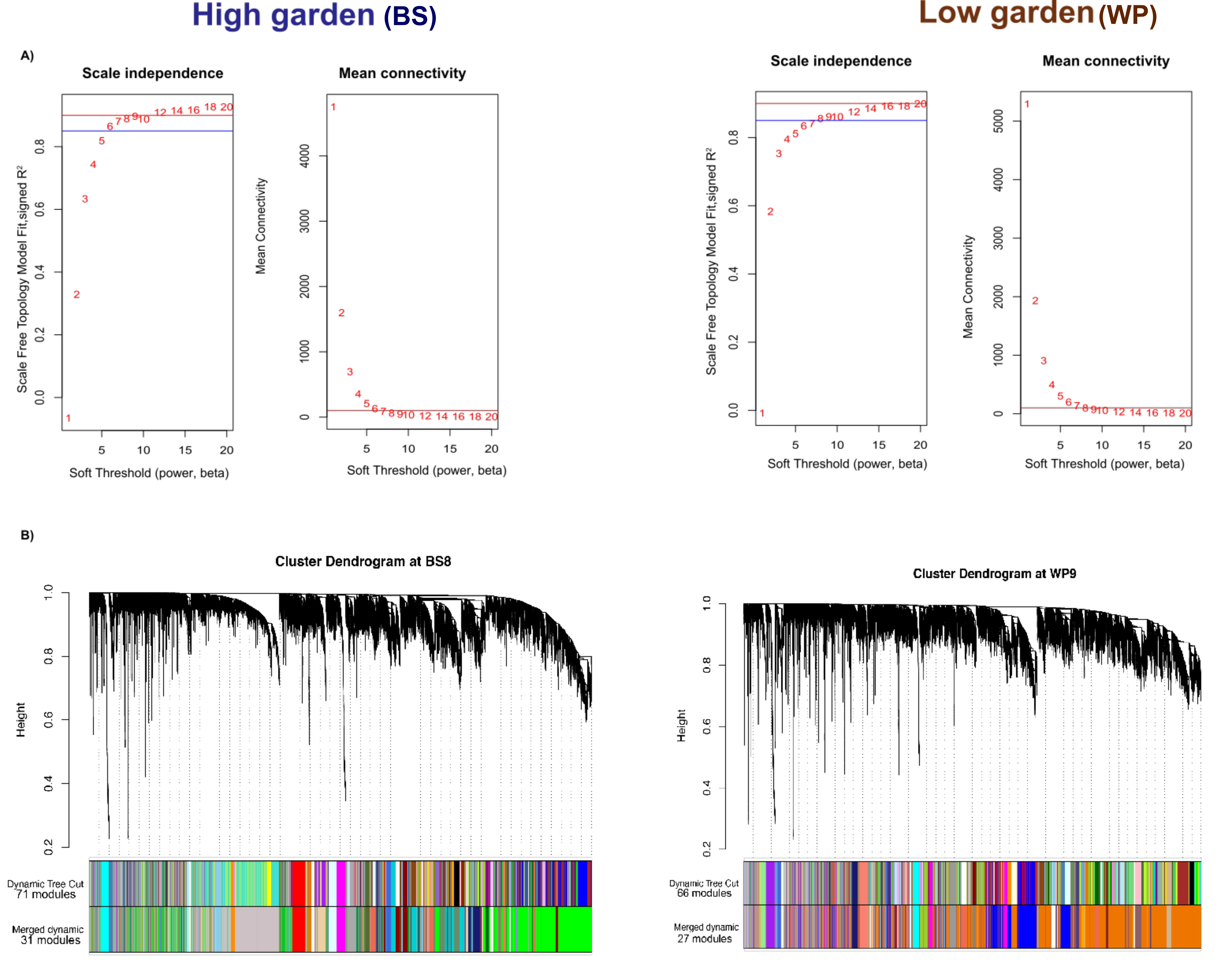
