## Supplemental Methods for "Variation in gene expression patterns across a conifer hybrid zone highlights the architecture of adaptive evolution under novel selective pressures"

**Supporting Methods S1: Details of the pipeline and summaries used for denovo transcriptome assembly and annotation.**

For species harbouring high levels of genetic diversity and large intergenic spaces, transcriptomes built by concatenating several individual reads can generate a fragmented assembly and underestimate heterozygosity since multiple reads across individuals will often be mapped to different transcripts (*pers. comm.* J. Wegrzyn). To avoid this issue, we built individual transcriptome assemblies using 20 samples representing each of the 10 populations and the two common gardens. These 20 *de novo* transcriptome assemblies were built using Trinity v.2.8.5 (Grabherr et al. 2011; Haas et al. 2013) with a *k*-mer length of 24 and a minimum contig size of 300bp. Redundancy reduction of this hyper-assembly was conducted through the tr2aacds pipeline in EvidentialGene v.2018.06.18 (Gilbert, 2013). The pipeline implemented within EvidentialGene first predicts coding regions and then removes transcripts that (a) are completely redundant, (b) exhibit high clustering similarity and (c) are perfect fragments of larger transcripts.

Transcripts that passed these steps were queried against the UniprotKB/Swissprot (release 2019_11) and PlantRef database (release 98) within Diamond (Buchfink et al. 2015) implemented within the EnTap pipeline (Hart et al. 2020) to filter out fungal and bacterial contamination from our reference transcriptome. Along with removal of contaminants EnTap was also used for annotating and obtaining gene ontology terms for the reference transcriptome. Through this we retained a total of 47,651 annotated and non-contaminated transcripts that were the starting point of further analyses. Read counts for each individual tree were extracted using RSEM with bowtie v.2.0 (Langmead & Salzberg, 2012) and ranged from 13 to 28 million reads per sample.

Additionally, we utilised a series of steps to assess the quality and the completeness of our transcriptome. First, we estimated the median length of contigs in our assembly (N50) using only the longest isoform per gene as well as the expression level sensitive matric that is considered suitable for RNAseq datasets called ExN50 (Haas et al. 2013). This process was done to evaluate the quality of our assembly and to explore the saturation of full length constructed transcripts as a function of the read depth. The N50 for our dataset was 654bp and Ex peak noted at E90 was 2kb corresponding to 13,872 genes. Second, the completeness of our assembly was assessed by using *Arabidopsis thaliana* as the reference database in the orthology-based algorithm implemented in BUSCO v.2.0 (Seppey et al. 2019). This approach identified 83.8% of the transcripts as matching and complete and 13% as missing. To evaluate the extent of conifer specific transcriptome space covered in our assembly we queried our transcripts against the publicly available *P. lambertiana* v.1.0 and *P. flexilis* (Liu et al. 2016) transcriptomes using blastp with an e-score threshold of 10^-50^ and 10^-100^, respectively. This process yielded 26,296 matches against the *P. lambertiana* transcriptome, covering 45% of the transcriptome space in *P. lambertiana*. For *P. flexilis* we identified 34,438 matches that covered 20% of the transcriptome space in *P. flexilis*. Third, we assessed mapping rates for individual fastq files to determine whether a sufficient amount of the transcriptome space was covered in the *de novo* assembled transcriptome. This process indicated an average mapping rate of 73% across 180 samples. Further, the mapping rate did not exhibit an upward bias towards the 20 samples that were used in the *de novo* assembly.

**Supporting methods S2: Pipeline details for SNP calling and filtering using ddRAD-seq genotyping.**

Processing of individual FASTQ files for the maternal trees was conducted using dDocent v 1.0 (Puritz et al. 2014). This process included the following steps implemented with dDocent a) read quality filtering with a PHRED score cutoff of 10, b) read grouping and a reference sequence assembly using the clustering algorithm CD-HIT with a within individual read coverage cutoff of 3 and an across individual read cutoff of 4, c) generating a reference assembly using the longest read in each cluster within a sequence similarity threshold of 0.89, d) read mapping to the *de novo* constructed reference assembly using BWA (default parameters except maximum mismatch value of 4 and a gap penalty of 6) and SNPs calling using FREEBAYES v 0.9.10 (Garrison & Marth, 2012).

Downstream processing of the resulting variant call format (VCF) file was performed using VCFTOOLS v.0.1.153 (Danecek et al. 2011) to remove indels, retain biallelic SNPs with at least 50% data and a minor allele frequency (MAF) cutoff of 0.003. Further processing involved the use of an absolute *F*_IS_ cutoff of 0.5 (keeping SNPs with values between 0.5 and −0.5), a minimum PHRED quality score of 20 and maximum depth per read set as the 75% percentile of depth distribution. These steps were performed using custom python scripts and yielded a total of 23,261 SNPs, which were used as the starting dataset for all subsequent analyses.

**Supporting methods S3: Details of steps and summary statistics for construction of the co-expression networks.**

Prior to constructing the networks at the level of maternal family for each garden, we filtered out transcripts below a median absolute deviation of 0.05. This resulted in a dataset of 22,757 transcripts at Bear Springs (BS) and 23,467 transcripts at White pockets (WP-Low garden) from which we constructed two separate co-expression networks in WGCNA v1.70 package (Langdelfer & Horvath, 2008) in R. For each garden we utilised the unsigned pearson’s adjacency matrix to identify a soft thresholding power (β) at which a scale-free topology can be attained. Biological networks are best represented by scale-free topology because they consist of few strongly connected hubs generating a network that is relatively stable to external perturbations. We allowed β to vary from 1 to 20 and picked a value at which R^2^ stabilises and mean connectivity remains relatively low (~100), following the WGCNA manual recommendation (Fig. S5.a). We proceeded with the final network construction using a β of 9 for WP and 8 for the BS. In both cases we were able to attain a nominal R^2^ of 0.85 or above and the mean connectivity dropped below 100.

Modules were identified following hierarchical clustering of the transcripts and a minimum size of 50 transcripts per module (Fig. S5.b). We subsequently merged very similar modules using a correlation coefficient of 0.70. These steps resulted in 31 modules in BS and 26 in WP which formed the basis of downstream analyses.
